# Intra-striatal AAV2.retro administration leads to extensive retrograde transport in the rhesus macaque brain: implications for disease modeling and therapeutic development

**DOI:** 10.1101/2020.01.17.910893

**Authors:** Alison R. Weiss, William A. Liguore, Jacqueline S. Domire, Dana Button, Jodi L. McBride

## Abstract

Recently, AAV2.retro, a new capsid variant capable of efficient retrograde transport in brain, was generated in mice using a directed evolution approach. However, it remains unclear to what degree transport will be recapitulated in the substantially larger and more complex nonhuman primate (NHP) brain. Here, we compared the biodistribution of AAV2.retro with its parent serotype, AAV2, in adult macaques following delivery into the caudate and putamen, brain regions which comprise the striatum. While AAV2 transduction was primarily limited to the injected brain regions, AAV2.retro transduced cells in the striatum and in dozens of cortical and subcortical regions with known striatal afferents. We then evaluated the capability of AAV2.retro to deliver disease-related gene cargo to biologically-relevant NHP brain circuits by packaging a fragment of human mutant *HTT*, the causative gene mutation in Huntington’s disease. Following intra-striatal delivery, pathological mHTT-positive protein aggregates were distributed widely among cognitive, motor, and limbic cortico-basal ganglia circuits. Together, these studies demonstrate strong retrograde transport of AAV2.retro in NHP brain, highlight its utility in developing novel NHP models of brain disease and suggest its potential for querying circuit function and delivering therapeutic genes in the brain, particularly where treating dysfunctional circuits, versus single brain regions, is warranted.

## INTRODUCTION

Adeno-associated viruses (AAVs) are small, non-enveloped viruses capable of packaging single-stranded DNA genomes up to ∼5 kb in length^1,2^. Originally discovered in 1965^3^, AAVs have become attractive agents for safely and effectively delivering gene cargo to a range of biological tissues (e.g. liver, muscle, retina, brain, kidney) in a wide variety of species (e.g. mouse, rat, cat, dog, pig, rabbit, horse, non-human primate, human)^1^. A number of AAV serotypes (e.g. AAV1-9, rh10, DJ, DJ/8) have been identified to date, as well as hundreds of naturally occurring capsid variants of each of these “parent” serotypes^4,5^. Differences between these AAVs are reflected in unique capsid structures, receptor specificity and tissue tropism^6^. For example, AAV6 and AAV8 transduce liver and skeletal muscle with high efficiency^4,7-9^, whereas AAV1, AAV2, AAV5 and AAV9 have been shown to transduce several types of cells in the CNS including neurons, astrocytes, and photoreceptors^2,10-13^.

Over the past decade, AAV-based gene therapies have begun to establish a track record of safety and success in human studies. For example, Luxturna (AAV2-RPE65) received Food and Drug Administration (FDA) approval in 2017 for the treatment of inherited retinal disease^14^ and Zolgensma (AAV9-SMN1) was FDA-approved in May of 2019 to treat spinal muscular atrophy type 1^15^. Additionally, there are ongoing early-stage clinical trials (recruitment and active phases) evaluating AAV-based gene therapies for neurological indications including, but not limited to, the treatment of Parkinson’s (AAV2-AADC and AAV2-GDNF), Huntington’s (AAV5-miHTT), Batten (AAV2-CUhCLN2, AAVrh10-CUhCLN2 AAV9-CLN3, AAV9-CLN6) and Alzheimer’s (AAVrh10-hAPOE2) diseases (*www.clinicaltrials.gov*). For most of these disorders, disease-specific pathology has been identified in multiple brain regions and so targeting AAV-mediated gene therapeutics to relevant circuits, versus single brain regions, would likely lead to a better therapeutic outcome for affected patients.

In addition to the immense therapeutic potential of AAVs for delivering disease-modifying constructs to the CNS, AAVs also have wide-ranging utility for neuroscience research from querying brain functions to creating animal models of disease. For example, several groups have downregulated or overexpressed genes in brain regions such as the hippocampus or prefrontal cortex to assess the impact on cognition, circuit dynamics, and synaptic plasticity^16-18^. More recently, AAVs have been used to deliver channel rhodopsins, or designer receptors, to localized brain regions, enabling researchers to stimulate/inhibit specific populations of neurons with optical signals (i.e. optogenetics^19^), or drugs (i.e. chemogenetics^20^). AAVs have also been used to create animal models of CNS disorders by delivering pathogenic constructs to brain regions of interest, such as the overexpression of α-synuclein in the striatum that results in progressive neurodegeneration and motor phenotypes characteristic of Parkinson’s disease^21-23^, or of expanded glutamine-encoding CAG repeats to create a striatal degeneration model of Huntington’s disease (HD)^21,24,25^, or of amyloid beta to reproduce features of Alzheimer’s disease ^26,27^.

Despite these recent successes, intra-parenchymal applications of AAVs in the CNS remain largely confined to the injected tissue region, or sub-region, because of restricted spread and limited capacity for anterograde and/or retrograde transport. One approach that has been used improve biodistribution of AAV constructs in the CNS is convection-enhanced delivery (CED), a technique that uses pressure-gradients to spread AAVs greater distances through brain tissue than simple diffusion alone^28-30^. Although this represents an improvement over conventional intraparenchymal infusion techniques, multiple injections are still needed to cover large brain regions or multiple structures. As such, the development of methods capable of distributing AAVs throughout multiple brain regions and biologically relevant circuits offers significant advantages both experimentally and clinically. One way to achieve this is with an AAV capsid capable of strong synaptic transport.

Recently, Tervo and colleagues generated a new AAV2 capsid variant capable of retrograde transport, named AAV2.retro, using an *in vivo* directed evolution approach in mice^31^. Mixed libraries of AAV *cap* variants were injected into discreet regions of the mouse CNS and variants were selected if they efficiently transported to neuronal cell bodies sending long-range projections to the site of AAV injection^31^. Since its creation, AAV2.retro has been used in mouse and rat models to target a multitude of CNS pathways including the amygdala via the ventral medial hypothalamus^32^, the thalamus via the anterior cingulate cortex^33^, the claustrum via the prefrontal cortex^34^, and more^35-37^. Taken together, these studies demonstrate that AAV2.retro is a powerful molecular tool capable of robust retrograde transport enabling the manipulation of neuronal pathways and circuits. However, it is unknown to what degree these features can be recapitulated in the larger and more complex primate brain. Therefore, we assessed the retrograde functionality of AAV2.retro in the nonhuman primate (NHP) brain by characterizing the biodistribution following stereotaxic injection of AAV2.retro expressing enhanced green fluorescent protein (AAV2.retro-eGFP) into the caudate and putamen of rhesus macaques, and comparing this to the biodistribution of its parent serotype, AAV2, injected into the same regions.

The ability to efficiently distribute AAV constructs throughout biologically relevant circuits in the brain offers significant advantages for the development of novel NHP models of neurological disease. Ongoing efforts in our laboratory are focused on creating an AAV-mediated model of Huntington’s disease (HD) via delivery of the disease-causing gene, mutant *HTT* (mHTT), into the caudate and putamen of adult rhesus macaques. Although it has been well established that the caudate and putamen are severely impacted in HD^38^, more recent studies have revealed that an extended network of structures throughout the cortex and basal ganglia are also affected^39,40^. Therefore, in order to refine our AAV-mediated NHP model to more closely mirror the widespread neuropathology documented in human HD patients, we further probed the capability of AAV2.retro to distribute a pathogenic fragment of mutant huntingtin protein (mHTT) throughout the rhesus macaque cortico-basal ganglia network.

## RESULTS

### Extensive retrograde transport in the rhesus macaque brain following MRI-guided intra-striatal delivery of AAV2.retro-eGFP

In order to investigate the retrograde transport capability of AAV2.retro in primate brain, naïve adult rhesus macaques were injected with AAV2.retro-eGFP bilaterally into the head of the caudate nucleus (80 ul at one injection site) and the putamen (150 ul over 2 injection sites spread apart by 4 mm). eGFP expression was driven from the human cytomegalovirus (CMV) promoter. The vector cartoon and surgical coordinates are illustrated in **Figure 1a** and **Table 1** summarizes each surgical case, including animal age, AAV construct, promoter, injectate titer/volume and post-surgical time to necropsy. Serum samples from all animals were tested for AAV2 neutralizing antibodies prior to surgery, and animals were selected only if they had less than 50% inhibition of transduction when serum was diluted to 1:20. There were no averse surgical events and all animals recovered fully post-infusion. Following a 4-week post-surgical interval, animals were euthanized, brains were collected and the biodistribution of AAV2.retro was visualized via immunohistochemical staining for eGFP in coronal tissue sections throughout the rostral to caudal extent of the brain. We observed dense eGFP positive (eGFP^+^) staining in the injected regions of the caudate (**Figure 1b**) and putamen (**Figure 1c**), with the spread partially filling each structure and the highest amount of transduction surrounding each site of injection. In the injected regions, the morphology of eGFP^+^ cells suggested that the majority of cells transduced were neurons, although transduced glia were noted as well but to a far lesser degree.

**Table 1.**
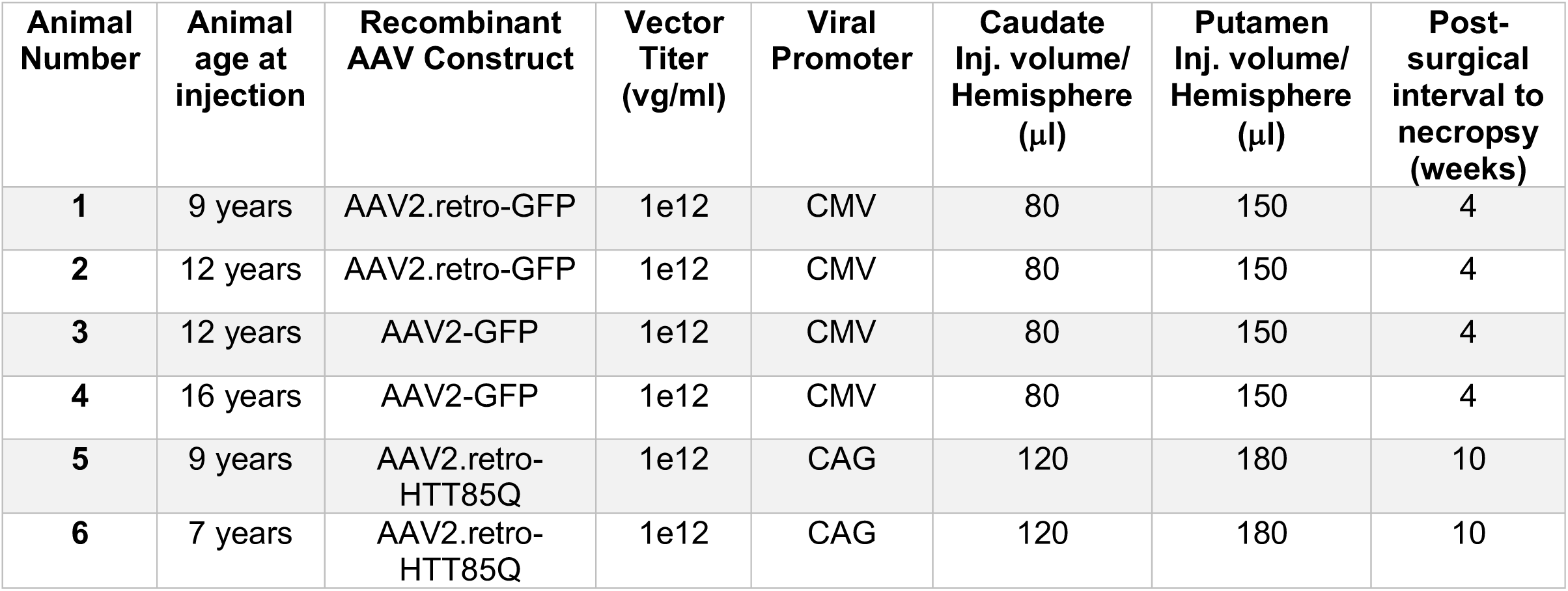
Summary of study participants and surgical cases. AAV- adeno-associated virus, vg- vector genomes, ml- milliliter, μl- microliter, GFP- green fluorescent protein, HTT85Q- mutant huntingtin protein bearing 85 CAG repeats, CMV- cytomegalovirus, CAG- chicken beta-actin promotor with a CMV enhancer element.

**Figure 1.**
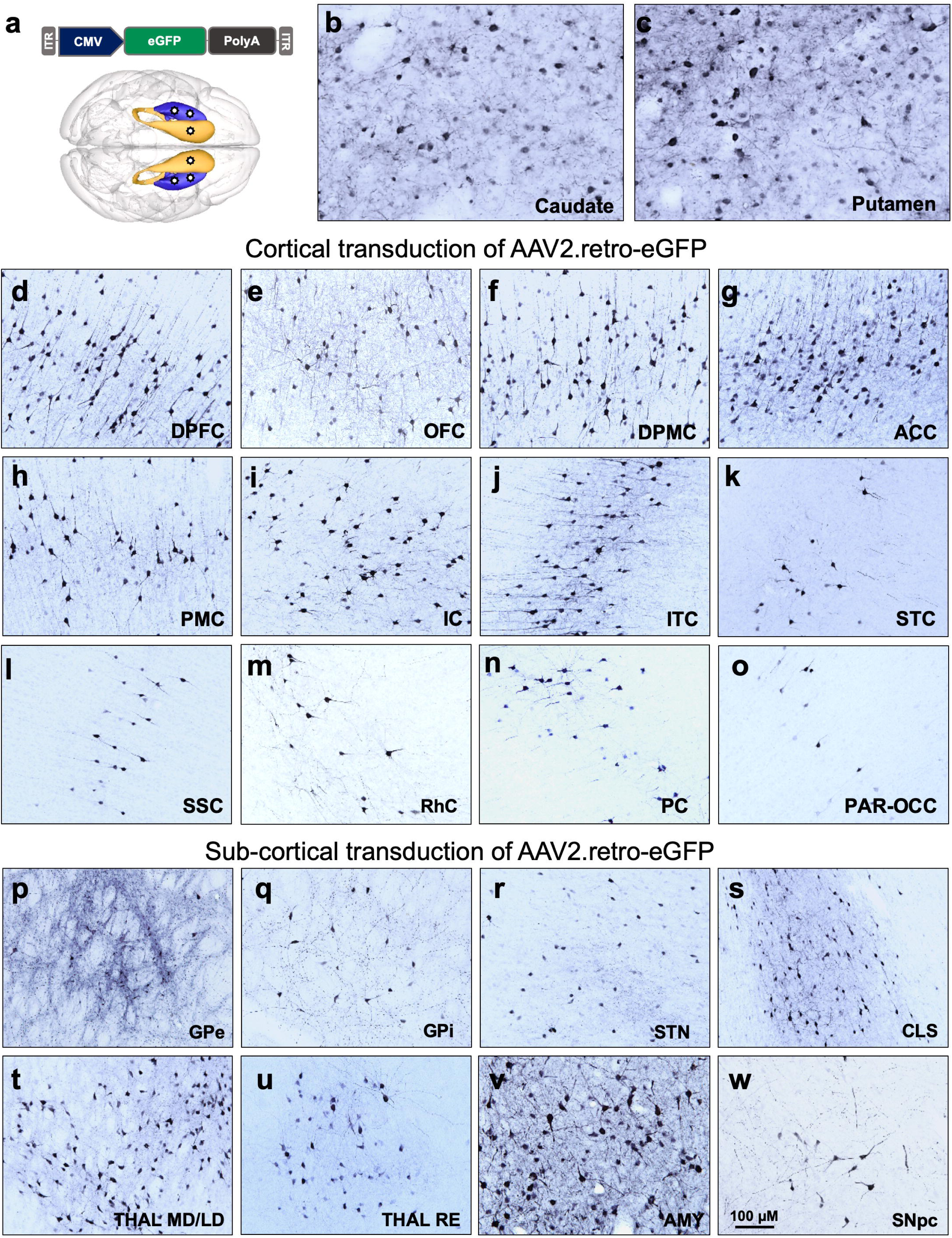
Biodistribution of AAV2.retro-eGFP following intra-striatal injection in adult rhesus macaques. (a) Illustrations of the AAV2.retro-eGFP vector construct and injection coordinates into the head of the caudate nucleus and the putamen. Robust eGFP expression in regions directly adjacent to injection sites of the caudate (b) and putamen (c). Following retrograde transport, additional eGFP expression was found in numerous cortical (d-o) regions and subcortical (p-w) structures. Abbreviations: ACC (anterior cingulate cortex), AMY (amygdala), CLS (claustrum), DPFC (dorsal prefrontal cortex, DPMC (dorsal premotor cortex), GFP (green fluorescent protein), GPe (globus pallidus, external), GPi (globus pallidus, internal), IC (insular cortex), ITC (inferior temporal cortex), OFC (orbitofrontal cortex), PAR-OCC (parieto-occipital cortex), PC (parietal cortex), PMC (premotor cortex), RHC (rhinal cortex), SNpc (substantia nigra pars compacta), SSC (somatosensory cortex), STC (superior temporal cortex), STN (subthalamic nucleus), THAL MD/LD (medial dorsal and lateral dorsal thalamic nuclei), THAL RE (reuniens nuclei). Scale bar = 100 microns.

In addition to the caudate and putamen, AAV2.retro injection resulted in widespread vector transport and eGFP expression in a total of sixteen cortical areas and eleven subcortical structures. Anatomical boundaries were defined using the rhesus macaque brain atlas developed by Saleem & Logothetis^41^. **Table 2** provides a comprehensive list of each of the observed transduced brain regions, and examples of a subset of these regions are depicted in **Figure 1**. Throughout most cortical regions, we observed dense staining of large pyramidal neurons in deep cortical layers (5/6) and lighter staining in neurons in the more superficial layers (2/3). Furthermore, the morphology of the eGFP^+^ cells that we observed in cortical areas were consistent with neurons, but not other cortical cell types like astrocytes or microglia. In the frontal lobe, robust eGFP^+^ expression was seen in the dorsolateral prefrontal cortex (DPFC, **Figure 1d**), orbitofrontal cortex (OFC, **Figure 1e**), dorsal premotor cortex (DPMC, **Figure 1f**), anterior cingulate cortex (ACC, **Figure 1g**) and primary motor cortex (PMC, **Figure 1h)**. In the parietal and temporal lobes, we observed eGFP^+^ staining in the insular cortex (IC, **Figure 1i**), the inferior and superior temporal cortices (ITC, **Figure 1j;** STC, **Figure 1k**), somatosensory cortex (SSC, **Figure 1l**) and the rhinal cortex (RhC, **Figure 1m**). Additionally, there was light eGFP staining in the parietal cortex (PPC, **Figure 1n**) and in the parieto-occipital cortical area (PAR-OCC, **Figure 1o**), but staining was largely absent in more posterior areas of the occipital cortex (not pictured).

**Table 2.**
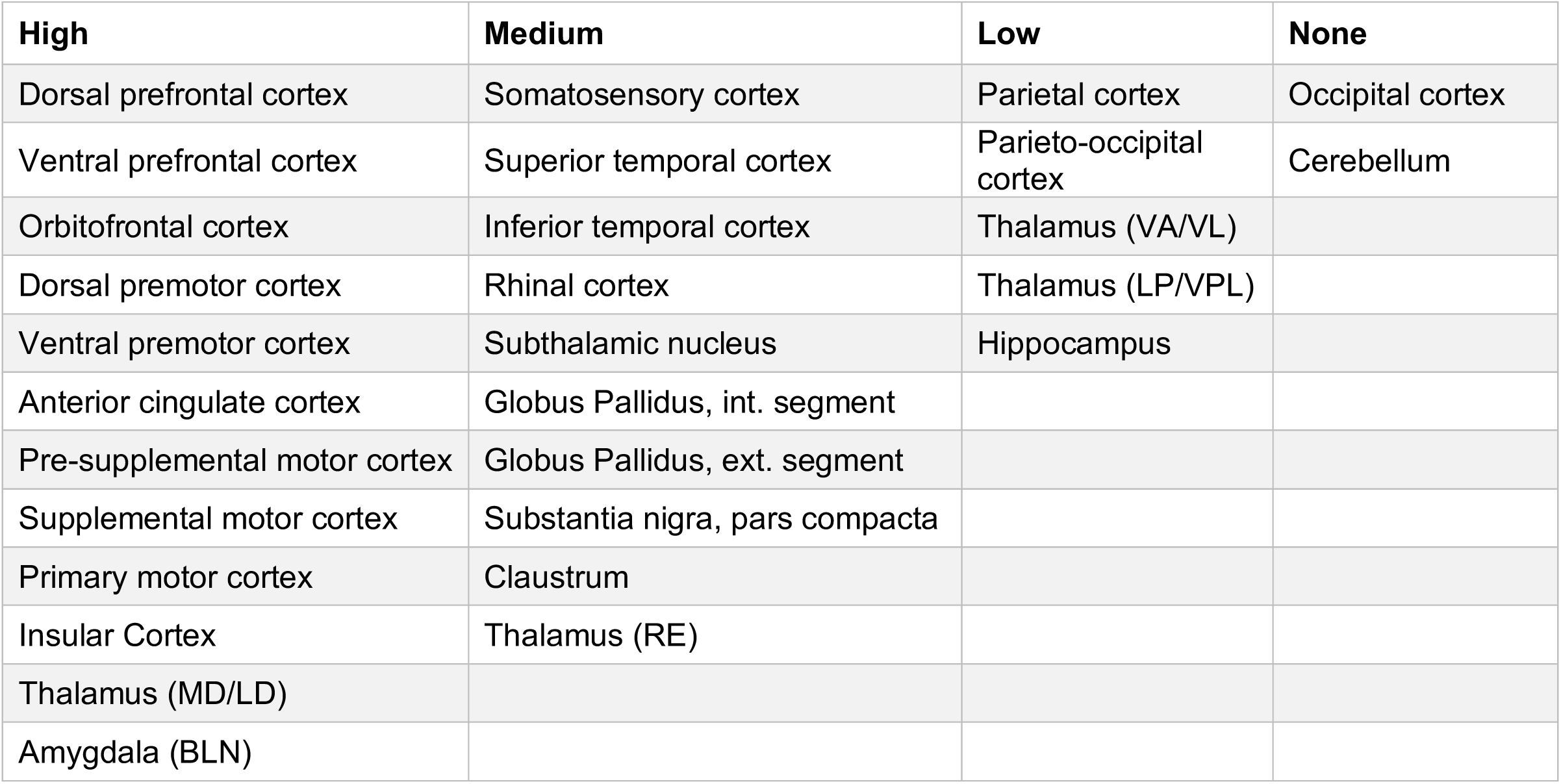
Relative levels of eGFP transgene expression in the rhesus macaque brain following intra-caudate and intra-putamen injection of AAV2.retro-eGFP. Brain regions were qualitatively ranked relative to one another, considering cell number and density, and placed into categories of high, medium, low and minimal/none. All regions listed here had less expression compared to the injected regions of the caudate and putamen. MD- medial dorsal, LD- lateral dorsal, RE- nucleus reuniens, BLN- basolateral nucleus, int.- internal, ext.- external, VA- ventral anterior, VL- ventral lateral, LP- lateral posterior, VPL- ventral posterolateral.

In addition to transduction of AAV2.retro in several cortical areas with known striatal afferent projections, we also detected AAV2.retro transduction in many subcortical structures as well. eGFP^+^ staining was seen in the external and internal globi pallidi (GPe, **Figure 1p**; GPi, **Figure 1q**), the subthalamic nucleus (STN, **Figure 1r**), the claustrum (CLS, **Figure 1s**), the medial dorsal and lateral dorsal nuclei of the thalamus (THAL MD/LD, **Figure 1t**), the reuniens nucleus of the thalamic midline group (THAL RE, **Figure 1u**), the basolateral nucleus of the amygdala (AMY, **Figure 1v**), and the substantia nigra pars compacta (SNpc, **Figure 1w**). The morphology of eGFP^+^ cells throughout these structures suggested that neurons were exclusively transduced, except in the medial dorsal region of the GPe and the medial region of the CLS, where we also observed a low number of eGFP^+^ cells with glial morphology, suggesting a small amount of local medial and lateral diffusion from the putamen to sub-regions of these two brain structures. Similarly, we postulate that the observed eGFP expression in the STN may have resulted from transport to this region resulting from primary transduction in the GPe.

### Comparisons of AAV2.retro-GFP and AAV2-GFP reveal significant differences in patterns of cortical and subcortical transduction

In order to further characterize and quantify the extent of retrograde transport by AAV2.retro, additional naïve adult rhesus macaques were injected with the parent serotype, AAV2, expressing eGFP. AAV2-eGFP was delivered into the same coordinates of the caudate nucleus and the putamen and at the same volume and titer (**Figure 1a, Table 1**). There were no averse surgical events and all animals recovered fully post-infusion. Following a 4-week post-surgical interval, brains were collected and the biodistribution of the AAV2-eGFP virus was visualized via immunohistochemical staining for eGFP in coronal sections. Cell counts of eGFP^+^ cells were manually collected from the regions of the caudate and putamen directly adjacent to the injection sites, as well as in 4 cortical areas (DPFC, DPMC, ACC, SSC) and in 2 subcortical structures (AMY and THAL). These cortical and subcortical regions of interest (ROIs) were selected due to their known dense connections with the striatum^42,43^ and relevance for future disease modeling and therapeutic development in our laboratory. **Figures 2a and 2b** illustrates panels of DAB immunohistochemistry staining in these brain regions from representative surgical cases and highlight a clear lack of AAV2 retrograde transport to extra-striatal regions (bottom row) compared to AAV2.retro (top row). A 2-way ANOVA was used to compare cell counts between serotypes (AAV2.retro, AAV2) and ROIs (Cd, Put, AMY, THAL, DPFC, DPMC, ACC, SSC). These analyses revealed significant main effects of serotype, F(1,18)=21.847 p=1.88E-4, and of ROI, F(8,18)=94.238 p=4.21E-13, but no interaction, F(8,18)=1.500 p=0.225. To further investigate these significant effects, we first conducted planned comparisons between serotypes for each ROI. **Figure 2c** illustrates these cell count comparisons separately for each ROI and serotype. Serotypes were compared for each ROI using one-tailed, independent-Samples t-tests. The results indicated that AAV2.retro-eGFP transduced significantly more cells than AAV2-eGFP in all of the extra-striatal ROIs: AMY (t(2)= 82.98, p<0.0001), THAL (t(2)=3.656, p=0.0337), DPFC (t(2)=79.66, p<0.0001), DPMC (t(2)=118.4, p<0.0001), ACC (t(2)=9.007, p=0.0061), SSC (t(2)=3.942, p=0.0294). On average, AAV2.retro-eGFP resulted in 7-fold higher levels of transduction in the putamen, and 3.5-fold higher transduction in the caudate, compared to extra-striatal ROIs.

**Figure 2.**
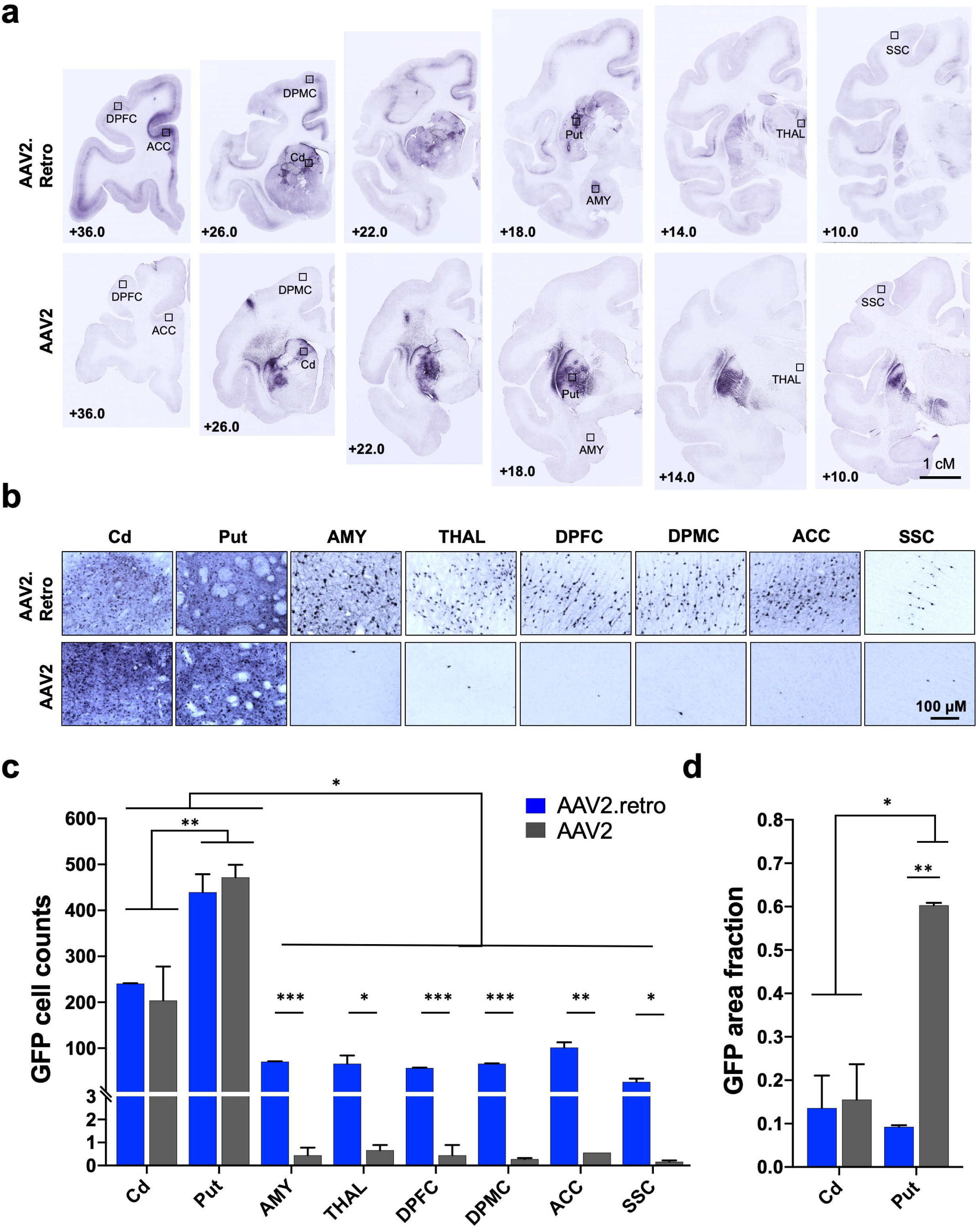
AAV2.retro retrograde transport is significantly higher compared to the parent serotype, AAV2. (a) Low and (b) high power photomicrographs of eGFP-stained, rhesus macaque coronal brain sections following injection of AAV2.retro-eGFP (top panel) or AAV2-eGFP (bottom panel) into the caudate and putamen. Rectangles in (a) illustrate the brain regions selected for high power photomicrographs displayed in (b). (c) Cell count analysis demonstrating significantly more GFP+ cells in the putamen compared to the caudate, irrespective of serotype, as well as significantly more GFP+ cells detected in extra-striatal regions in animals injected with AAV2.retro-eGFP compared to AAV2-eGFP. (d) Area fraction fractionator analysis showing a significantly higher area fraction of GFP positivity in the putamen in animals injected with AAV2-eGFP compared to AAV2.retro-eGFP. Error bars in both graphs represent SEM. Abbreviations: ACC (anterior cingulate cortex), AMY (amygdala), Cd (caudate), DPFC (dorsal prefrontal cortex, DPMC (dorsal premotor cortex), Put (putamen), SSC (somatosensory cortex), THAL (thalamus). * p<0.05, ** p<0.01, *** p<0.001. Scale bar 2a = 1cm, Scale bar 2b=100 microns.

Next, we compared cell counts between ROIs for each serotype separately by one-way ANOVA with planned comparisons using one-tailed independent sample t-tests. The results indicated that the number of eGFP^+^ cells differed significantly depending on brain region for both AAV2.retro (F(8,9)=73.936, p<0.0001) and AAV2 (F(8,9)=38.639, p<0.0001). Post-hoc comparisons revealed significantly higher cell counts in the putamen than in the caudate for both AAV2.retro (p=0.0003) and AAV2 (p=0.0018), and significantly higher cell counts in both of the injected regions than any of the other regions for AAV2 and AAV2.retro (all p-values<0.05). The higher cell count in the putamen versus the caudate is intuitive given the substantially larger volume of vector injected into the putamen (150 ul) compared to the caudate (80 ul) for both serotypes. There were no significant differences in cell counts between any of the other regions for both serotypes (all p-values n.s.). The results of these comparisons are reported in detail in **Supplemental Table S1**.

Additionally, there were no significant differences in cell counts between AAV2 and AAV2.retro in the caudate (t(2)=0.497, p=0.334) or putamen (t(2)=-0.680, p=0.283), suggesting similar levels of transduction between serotypes in the regions directly adjacent to the injection sites. Interestingly, even though the cell counts were similar directly adjacent to the injection sites, we observed robust differences between AAV2 and AAV2.retro in the amount of spread within the putamen from the site of injection. In particular, AAV2 spread throughout the putamen to a far greater extent than AAV2.retro, resulting in much larger regions of transduction clearly visible in multiple coronal sections, **Figure 2a**, bottom row. To assess these serotype-based differences, we quantified the spread of each vector in the injected regions of the caudate and putamen using the Area Fraction Fractionator tool (MBF Bioscience), calculating the area of eGFP^+^ cells per the area of each structure. **Figure 2d** illustrates these measurements, which were compared using a 2-way ANOVA (Serotype x ROI). The results revealed a significant Serotype*ROI interaction, F(1,4)=19.57, p=0.012, as well as significant main effects of Serotype (F(1,4)=22.85, p=0.009), and ROI (F(1,4)=13.30, p=0.022). Planned comparisons using one-tailed t-tests indicated that the area of transduction was significantly larger in the AAV2-injected putamen than the AAV2.retro-injected putamen, t(2)=75.37, p<0.0001, whereas the area of transduction did not significantly differ between AAV2-injected caudate and AAV2.retro-injected caudate, t(2)=0.18, p=0.437. There were additional differences such that AAV2 spread over a larger area of the putamen than the caudate, t(2)=5.492, p=0.016, whereas AAV2.retro had similar spread in both of the injected regions t(2)=0.5726, p=0.312.

As described above, AAV2.retro-eGFP injections resulted in eGFP^+^ staining in many cortical and subcortical structures in addition to the eight regions used for the cell counting analysis. Since brain-wide manual cell counting is beyond the scope of this study, we performed a qualitative ranking to assess eGFP staining density in each of the extra-striatal brain regions where staining was found, relative to one another. Regions were rated as having either high, mid or low-level staining density (or no staining) and ratings are reported in **Table 2.** To depict brain-wide differences between the two serotypes, and to highlight the pattern of AAV2.retro biodistribution in the rhesus macaque brain following intra-striatal injection, the ranked transduction patterns for each serotype are illustrated in a cartoon brain graphic generated using an in-house rhesus macaque T2-weighted MRI template/atlas to define each transduced region and then visualized graphically using paraView 5.7.0 software (**Figure 3**) ^44^. High-level transduced brain regions are depicted in dark blue, mid-level regions are depicted in royal blue, lower-level regions are depicted in turquoise, and regions with minimal to no staining are shown in gray. Injected ROIs are depicted in yellow for reference. AAV2.retro transduction largely followed an anterior to posterior gradient, with the highest levels of eGFP expression in the frontal lobe (pre-frontal and motor areas) and mid to low level staining in the temporal and parietal lobes, respectively. No discernable eGFP staining was detected in the occipital lobe nor the cerebellum.

**Figure 3.**
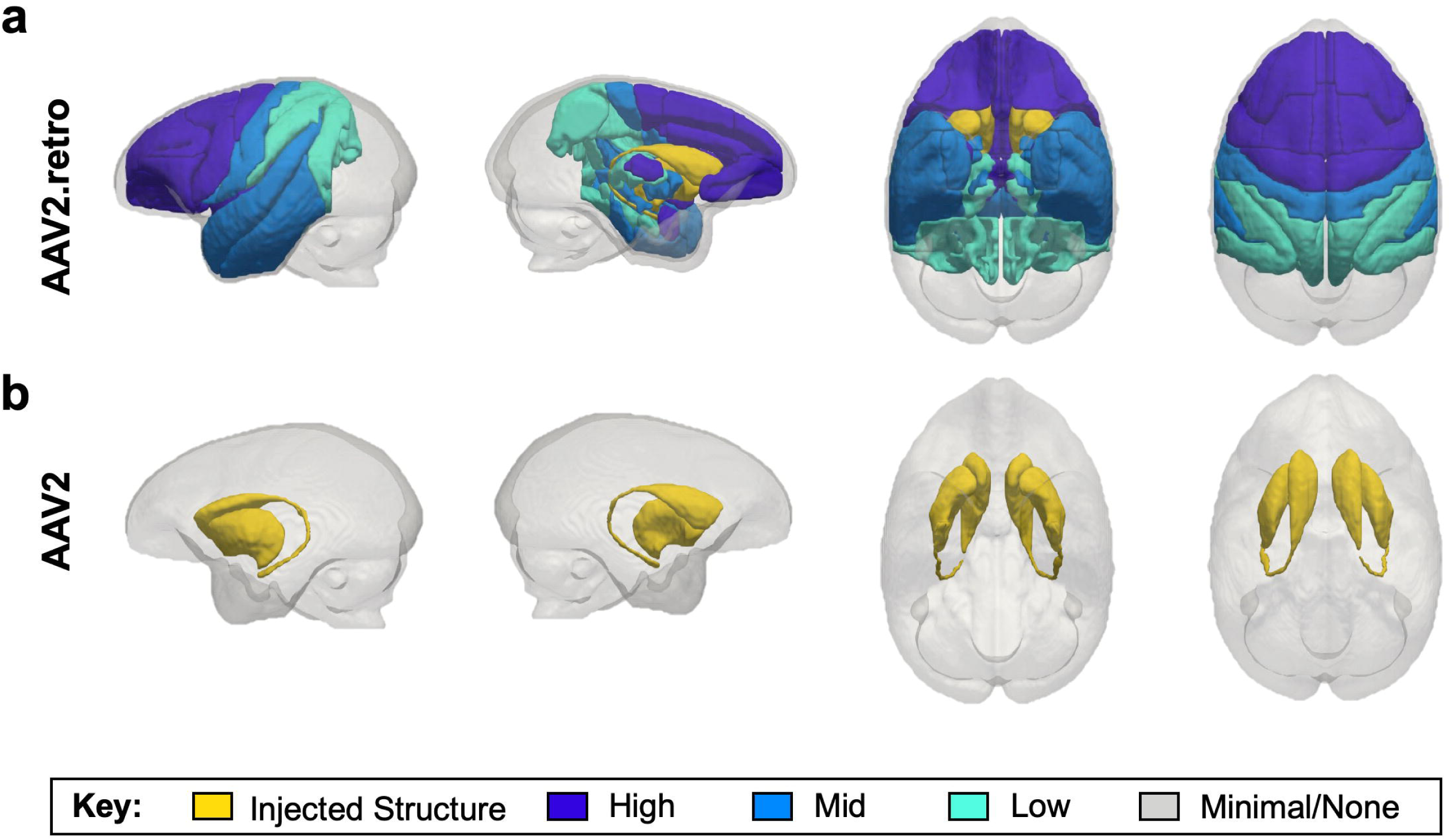
Cartoon brain graphic illustrating the brain-wide transduction patterns for AAV2.retro and AAV2. The ‘glass brain’ graphic was generated from an in-house rhesus macaque T2-weighted MRI template and visualized using paraView 5.7.0 software^44^. Levels of transduction were ranked based on eGFP expression in each brain region, relative to one another. High-level transduced brain regions are depicted in dark blue, mid-level regions are depicted in royal blue, lower-level regions are depicted in turquoise, and regions with minimal to no staining are shown in gray. Injected ROIs are depicted in yellow for reference. (a) The density of AAV2.retro transduction largely followed an anterior to posterior gradient, with the highest levels of staining in the frontal lobe (pre-frontal and motor areas) and mid to low level staining in the temporal and parietal lobes, respectively. No discernable staining was detected in the occipital lobe nor the cerebellum. (b) AAV2 transduction was restricted to the injected regions only.

### Intra-striatal injection of AAV2.retro induces mHTT expression distributed throughout multiple functional circuits in the cortico-basal ganglia network

To evaluate the capability of AAV2.retro to deliver disease-relevant cargo to biologically relevant circuits in the NHP brain, naïve adult rhesus macaques were injected with AAV2.retro expressing a pathogenic fragment of mutant huntingtin protein (mHTT) bearing 85 CAG repeats (AAV2.retro-HTT85Q) into the head of the caudate and the putamen. mHTT85Q expression was driven from a chicken beta-actin promoter with a CMV early enhancer element (CAG promoter). See **Figure 4a** for vector diagram and **Figure 4b** for a cartoon of the unilateral surgical injection sites. To improve localized spread within the injected regions, we increased the number of injection sites in the caudate to 2 (one pre-commissural; one post-commissural) and increased the volume of injection for each structure (see **Table 1** for summary of surgical parameters). Additionally, we increased the post-surgical interval from 4- to 10-weeks post-surgery to allow enough time for the formation of pathological mHTT protein aggregates. There were no averse surgical events and all animals recovered fully post-infusion. At 10-weeks post-surgery, brains were collected and the biodistribution of intracellular mHTT was visualized in coronal sections via immunohistochemical staining using the mHTT 1-82aa antibody (1-82aa). We observed 1-82aa^+^ staining throughout the head of the caudate (**Figure 4c**) and putamen (**Figure 4d**), confirming the formation of aggregated intracellular mHTT aggregates in these structures. We also observed 1-82aa^+^ staining in dozens of cortical regions and subcortical structures, with examples shown here including the AMY (**Figure 4e**), the THAL (**Figure 4f**), the DPFC (**Figure 4g**), the DPMC (**Fgure 4h**), the ACC (**Figure 4i**), and the SSC (**Figure 4j**). See **Supplemental Table S2** for summary of all the regions in which we detected the formation of mHTT 1-82aa^+^ aggregates. In general, the brain-wide distribution of intracellular aggregates largely mirrored that of the eGFP^+^ staining that we described in the earlier cases reported here, with a few limited exceptions. Notably, we observed dense mHTT aggregates in the STN and claustrum, regions in which we observed somewhat lower relative levels of eGFP transduction. These variations could be due to differences in the volumes and locations of the eGFP and HTT85Q injections, and/or the result of retrograde transport from regions outside of the striatum that were reached by diffusion from the injection sites, such as the GPe. We stained coronal brain sections using the microglial antibody ionized calcium-binding adapter molecule 1 (Iba1) to investigate a potential immune response to the AAV2.retro vector or to the mHTT transgene expression. We found activated Iba1^+^ microglia in the needle tracts of the caudate and putamen; however, no diffuse microgliosis was seen throughout the injected brain regions nor in any of the striatal afferent brain regions expressing mHTT. Similarly, inspection of neuronal nuclei (NeuN)-stained tissue showed no loss of neurons in any brain regions.

**Figure 4.**
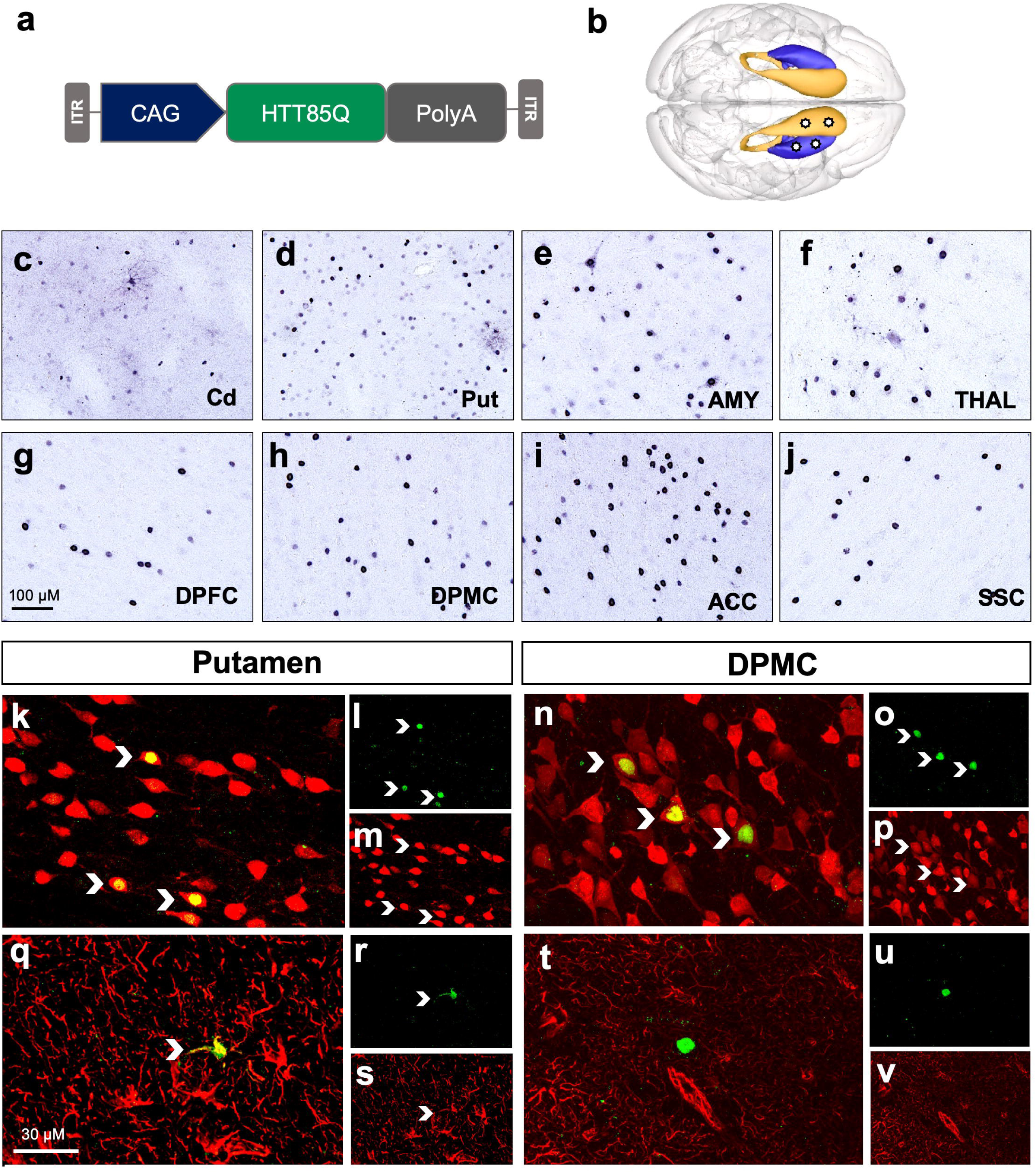
AAV2.retro delivery of mutant mHTT into the rhesus macaque striatum leads to aggregate formation in disease-relevant brain regions. (a) AAV2.retro vector cartoon and (b) surgical injection graphic depicting unilateral injection sites of AAV2.retro-HTT85Q into the caudate and putamen. (b) 1-82aa staining of mHTT protein showing mHTT^+^ aggregates in the caudate (c), putamen (d) and several cortical and subcortical brain regions. Examples shown here include the AMY (e), THAL (f), DPFC (g), DPMC (h), ACC (i) and SSC (j). Double label immunofluorescence of HTT 1-82aa/NeuN (k-p) and HTT 1-82aa/GFAP (q-v) in the injection site (putamen) and a distal brain region (DMPC). Transduced neurons and astrocytes are indicated with chevrons. Abbreviations: ACC (anterior cingulate cortex), AMY (amygdala), Cd (caudate), DPFC (dorsal prefrontal cortex, DPMC (dorsal premotor cortex), Put (putamen), SSC (somatosensory cortex), THAL (thalamus). Scale bar in g=100microns, scale bar q=30 microns.

To investigate the cell-type specificity of mHTT expression, we next compared the injected regions of the caudate and putamen versus the transduced striatal afferents using double-label immunofluorescence. The examples shown in **Figure 4k-v** are from the putamen (left panel) and the DPMC (right panel), but are representative of the other injected region as well as the other striatal afferents that also exhibited mHTT expression. Co-localization of HTT1-82aa^+^/NeuN^+^ staining was used to identify transduced neurons, and HTT1-82aa^+^/GFAP^+^ staining was used to identify transduced astrocytes. In the putamen, we detected co-localization of mHTT with both neurons (**Figure 4, k-p)** as well as astrocytes **(Figure 4 q-v)**, although astrocyte transduction was far less prevalent. In contrast, we only detected transduced neurons in striatal afferents including the DPMC (**Figure 4 n-p**) but did not detect examples of transduced astrocytes (**Figure 4, t-v**), These data suggest that AAV2.retro was transported to these distal regions through uptake at the nerve terminal in the striatum and transported back to the cell body, rather than via long-range diffusion from the injection sites.

Broadly, the observed pattern of AAV2.retro-HTT85Q biodistribution resulted in HD-pathology throughout the same circuits of the cortico-basal ganglia network that were targeting via AAV2.retro-eGFP delivery into the caudate and putamen. Moreover, the pattern of transgene expression shown here models the distribution of mHTT protein and aggregate formation seen in patients with HD, where cognitive, sensorimotor and limbic cortico-basal ganglia circuits are each affected, leading to the wide array of complex symptoms experienced by patients. A schematic summarizing each brain region we found transduced by AAV2.retro following intra-striatal delivery, and then placed into the framework of functional cortico-basal ganglia circuits, is shown in **Figure 5.** In brief, we observed transduction in cognitive circuits that include the dorsal and ventral prefrontal cortices, rhinal cortex and the hippocampus; in sensorimotor circuits that include premotor, supplemental motor, primary motor and somatosensory cortices, and in limbic circuits that include the temporal, cingulate, and orbitofrontal cortices and the amygdala. In addition, we observed AAV2.retro transduction in regions of the basal ganglia and thalamus that were presumably reached by our vector via their afferent projections to the striatum (SNpc, THAL) and/or by diffusion/transport from the nearby injection sites (GPe, STN). Taken together, these results highlight that the retrograde capabilities of AA2.retro enable the delivery of viral constructs simultaneously to dozens of brain regions comprising multiple functional brain circuits from injections into two focal brain regions.

**Figure 5.**
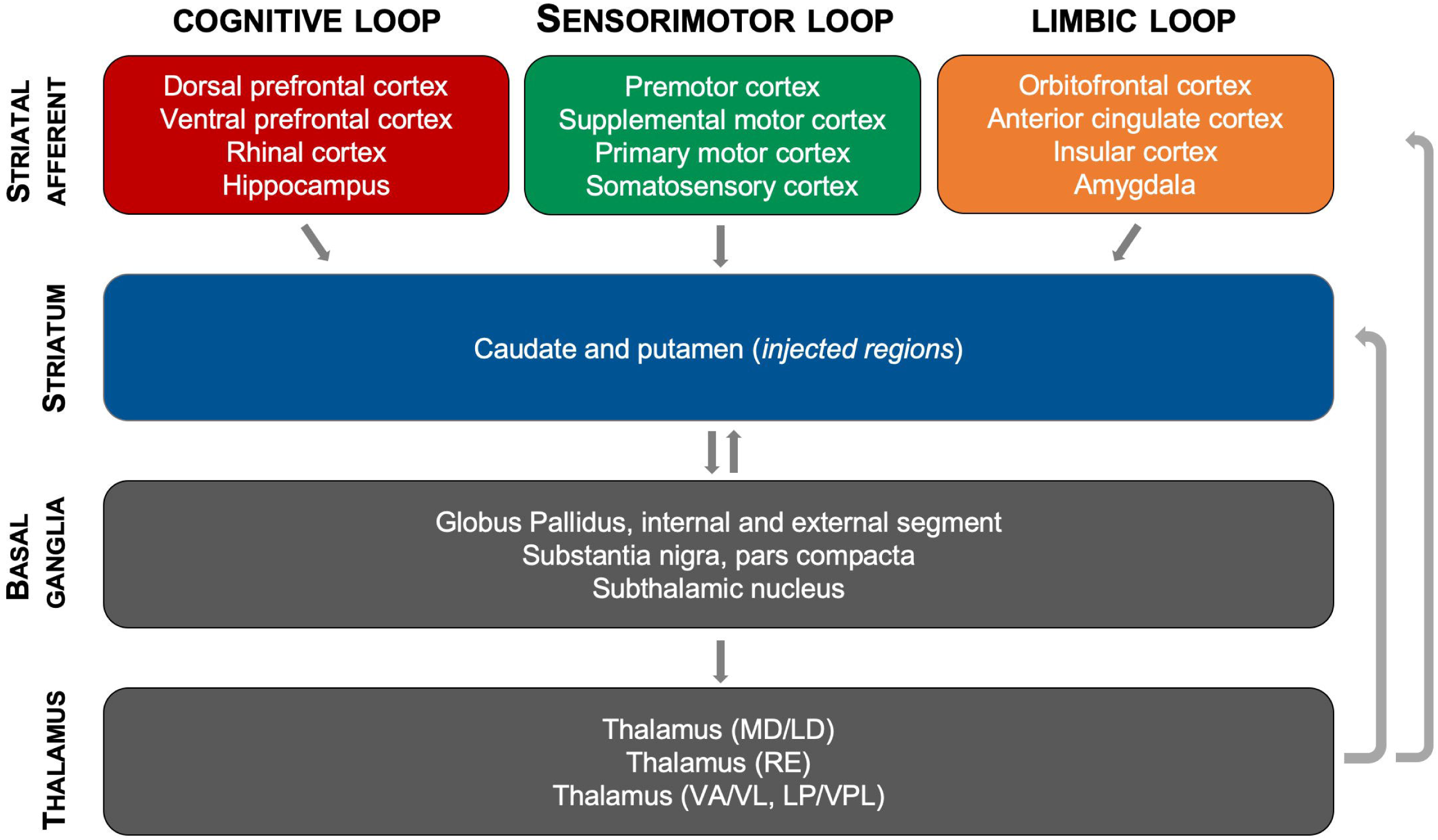
Schematic of the functional cortico-basal ganglia circuits targeted via intra-striatal delivery of AAV2.retro. Delivery of AAV2.retro into the caudate and putamen lead to transgene expression (both eGFP and mHTT85Q) in dozens of highly interconnected brain regions comprising cognitive, sensorimotor and limbic cortico basal ganglia functional neural networks.

## DISCUSSION

The results reported here provide clear evidence of extensive retrograde transport in the rhesus macaque brain by a newly derived AAV2 capsid variant, AAV2.retro^31^. The brain-wide biodistribution pattern of AAV2.retro following intrastriatal injection was significantly more widespread than that of its parent serotype, AAV2, with AAV2.retro transducing cells throughout the cortico-basal ganglia network and AAV2 transducing cells limited to the targeted regions of the caudate and putamen. The findings here in adult rhesus macaques closely recapitulate those previously reported in adult mice following intra-striatal delivery of similar AAV2.retro constructs^31^, and indicate that the mechanism of retrograde transport used by AAV2.retro is conserved across species.

We observed clear serotype differences of vector spread in the region of injection, with AAV2 transducing a significantly larger area of the putamen compared to AAV2.retro. A likely interpretation for this finding is that, because such a large proportion of the infused AAV2.retro viral particles were taken up at the injection sites by nerve terminals and transported to distal brain regions, fewer AAV2.retro viral particles were available to diffuse into the surrounding regions of the caudate and putamen. Future studies employing these vectors to target multiple brain structures or circuits might be able to take advantage of each of these features by combining both serotypes into a single bolus infusion in order to maximally transduce the area of injection with AAV2, yet also benefit from the enhanced retrograde functionality of AAV2.retro.

It is noteworthy that other AAV serotypes exhibit low to moderate levels of retrograde transport, such as AAV1^45,46^ and AAV6^47^, but by comparison the degree of retrograde transport is greatly enhanced in AAV2.retro^31^. There are, however, other AAV capsid variants that have been recently reported to also transport robustly in the NHP brain, such as AAV-HBKO^48^. Naidoo and colleagues demonstrated that intra-thalamic delivery of AAV-HBKO expressing eGFP resulted in robust anterograde transport to deep layers of many cortical regions with thalamic efferents including the ACC, DPFC, VPFC, DPMC, VPMC, PMC, SSC, and ITC. The authors also report evidence of moderate retrograde transport to some subcortical structures and to regions of the brain stem and spinal cord^48^. This contrasts with the pattern of nearly exclusive retrograde transport seen here with AAV2.retro as well as transduction of both deep and superficial layers of cortex. Our findings reflect the known cortical afferent inputs into the caudate and putamen which have been previously characterized in tracer studies showing innervation from both deep and superficial layers. Together, these two studies illustrate the feasibility of using novel, designer AAV serotypes with a high propensity for axonal transport to distribute genetic cargo throughout brain circuits by targeting specific structures with a high degree of connectivity in the CNS, such as the thalamus and the striatum. For example, in the current study we demonstrated that AAV2.retro efficiently transduces cognitive, sensorimotor, and limbic cortico-basal ganglia circuitry via intra-striatal delivery, and the work by Naidoo and colleagues with AAV-HBKO demonstrated biodistribution throughout a different network of prefrontal, motor, temporal, and subcortical regions following thalamic infusion^48,49^. Taken together, these studies illustrate how widespread transduction patterns can be achieved even in the large NHP brain by selecting a serotype with high propensity for transport and by selectively choosing the site of injection to target specific populations of neuronal afferents/efferents depending on the network of interest or the specific disease being targeted.

Additional work has explored the merits of more systemic administration routes with the potential to increase brain-wide AAV biodistribution. One approach has been to employ capsids capable of crossing the blood brain barrier (BBB), such as AAV9^50^ and AAV.PHPb^51^. Work with these capsids indicates that they transduce broad regions of the CNS when administered intravenously (IV), but with relatively low efficiency compared to intra-parenchymal delivery^50,51,52^. Recent work from our group shows that intra-CSF administration of AAV-PHP.b leads to greater neuronal and astrocytic transduction compared to intravascular infusion. However, transduction was still significantly greater in cortical areas compared to subcortical brain regions^52^. In contrast, intraparenchymal delivery of vectors like AAV2.retro and AAV-HBKO can target widespread brain regions that encompass both cortical and subcortical structures. Hence, these and other new capsid variants capable of enhanced axonal transport may offer a method to target widespread neuronal circuits with greater efficiency and selectivity. The downside to this strategy compared to intra-IV and intra-CSF delivery techniques, however, is a more complex and invasive surgery.

With the advancement of neuroimaging techniques, it has become apparent that many neurological diseases are characterized by dysfunction on the level of brain networks, rather than individual structures. For example, although in HD the caudate and putamen are profoundly affected, severe dysfunction is also apparent in other basal ganglia structures and in several motor, cognitive, and limbic cortical areas as well^38-40,53^. Likewise, the hippocampus is known to be particularly vulnerable in Alzheimer’s Disease, but atrophy also occurs in brain regions throughout the temporal cortex, the prefrontal cortex, the parietal cortex, and the thalamus^54,55^. Therefore, an important limitation to current gene-therapy approaches for these disorders are that they do not target the entire dysfunctional network. Although the approaches targeting single brain regions are predicted to offer some therapeutic benefit to patients based on pre-clinical animal research, it is logical to postulate that treating the entire dysfunctional network, rather than just one structure, could result in greater therapeutic benefit.

In summary, the studies reported here show that AAV2.retro exhibits robust retrograde transport when injected into the NHP striatum, a phenomenon that was first demonstrated using a control eGFP transgene and then verified by expressing the disease-causing gene, mHTT, as a proof of concept. The transport and widespread gene delivery were well-tolerated and did not lead to adverse pathological events nor neurological symptoms in any animals, at least out to the four- and ten-week study timelines conducted here. Efforts to characterize the structural and functional consequences of AAV2.retro-mediated delivery of mHTT to these cortico-basal ganglia circuits in a larger cohort of rhesus macaques over a longer timeframe are ongoing. The dearth of genetically- and biologically relevant large animal models of disease has been identified as a significant limitation hampering the translation of new scientific discoveries to the clinic. There is an immense cost and effort required to generate and maintain lines of transgenic NHPs and the work presented here represents some the first to test the potential for this new AAV serotype to become a tool for the rapid generation of novel NHP models of neurodegenerative disease, something for which there is great interest in the biomedical research community. Additionally, our findings offer a solid framework for future studies in large animal models and human patients using AAV2.retro as a gene therapy tool to deliver therapeutics for myriad brain disorders, particularly those where targeting multiple brain regions and/or circuits would result in a better clinical outcome for the patient including HD, Parkinson’s disease, Alzheimer’s disease, amyotrophic lateral sclerosis, lysosomal storage diseases, among others.

## MATERIALS AND METHODS

### SUBJECTS

Six adult female rhesus macaques were involved in this study (age 7-12). Animals were selected based upon their negative neutralizing antibody status for AAV2 virus. Monkeys were pair housed on a 12-hour light/dark cycle lighting schedule, provided with monkey chow rations BID, and given ad libitum access to water. Macaques were observed daily by trained veterinary technicians, and body weights of all animals were recorded monthly. All experimental procedures were approved by the Institutional Animal Care and Use Committee and the Institutional Biosafety Committee at the Oregon National Primate Research Center and Oregon Health and Science University. All guidelines specified in the National Institutes of Health Guide for the Care and Use of Laboratory Animals were strictly followed.

### VECTOR PREPARATIONS

Recombinant AAV vectors were generated by a scalable co-transfection procedure in the OHSU/ONPRC Molecular Virology Support Core. pAAV2.retro capsid plasmids were kindly provided by the Karpova lab at the Howard Hughes Medical Institute (HHMI), Janelia Research Campus via a Material Transfer Agreement with OHSU. Plasmids containing the sequence for the N171 N-terminal fragment of human HTT bearing 85 CAG repeats were manufactured at Genscript and subsequently cloned into a transgene cassette flanked by viral ITRs. pAAV2 capsid plasmids were supplied by the OHSU/ONPRC Molecular Virology Support Core. Viral vector preparations were prepared as described previously^52^. Briefly, mammalian HEK293 producer cells were transfected with plasmids carrying the transgene cassette flanked by viral ITRs (sspAAV-CMV-eGFP, sspAAV-CAG-HTT85Q), a rep-cap expression construct encoding the sequence for the AAV2.retro or AAV2serotype capsid, and a helper plasmid expressing adenoviral E2a, VA, and E4-orf6. Viral lysates were treated with Benzonase to remove residual plasmid, and virus was purified over an Iodixanol step gradient. Gradient fractions containing intact virus and excluding empty particles were harvested, and the final virus preparation was buffer-exchanged into DPBS+5% glycerol+35mM NaCl. Quality control was performed to ensure purity by viral capsid protein evaluation with silver staining on SDS-PAGE. Viral titers for AAV2.retro.CAG.85Q were determined by quantitative PCR of purified vector particles using a CAG primer/probe set: Forward: 5’-CCATCGCTGCACAAAATAATTAAAA, Reverse: 5’-CCACGTTCTGCTTCACTCTC-3, Probe: 5’-CCCCTCCCCACCCCCAATTTT. Viral titers for AAV2.CMV.eGFP were determined by quantitative PCR of purified vector particles using a viral ITR primer/probe set: Forward: 5’-GGAACCCCTAGTGATGGAGTT, Reverse: 5’-CGGCCTCAGTGAGCGA, Probe: 5’-CACTCCCTCTCTGCGCGCTCG.

### NEUTRALIZING ANTIBODY ASSAY

Whole blood was collected in Vacutainer Serum Collection Tubes (BD) and serum was collected following centrifugation at 2500 rpm for 10 min and stored at -80C until analysis. Neutralizing antibody screening assays were carried out as previously described^52^. Briefly, assays were performed in 96 well format with 5 x 10^4 CHO-Lec2 cells per well. Serial dilutions of study participant serum were pre-incubated with 1E+9GC of AAV2 reporter virus for 1 hour at 37°C and then added to cells that were infected with Adeno Helper virus. After 48h, Promega Bright-Glo substrate was added to the cells and luciferase expression was quantified using the Biotek Synergy Mx luminometer. Both positive and negative monkey sera controls were included with each assay. All animals selected for the study had less than 50% inhibition of transduction when serum was diluted to 1:20.

### NEUROIMAGING

On the day of surgery, all monkeys received T1 MRI scans to determine coordinates for the AAV injections. Anesthesia was first induced with Ketamine HCl (10 mg/kg IM), followed by maintenance anesthesia with isoflurane gas vaporized in 100% oxygen. Animals were placed in a Crist NHP-specific stereotaxic frame in which they remained for the duration of the scan and subsequent surgery. A twelve-minute T1-weighted scan was collected on a Siemens Prisma scanner using a surface coil. Brain images were subsequently examined using Osirix software and the injection coordinates were selected and transferred from MRI-space to stereotaxic space.

### STEREOTAXIC SURGERY

Monkeys were taken directly from the MRI to the operating room immediately following their scan. Pre-operative care consisted of overnight food restriction (surgery SOP) for 12 hours and general examination to ensure that patient health was adequate to withstand the procedure. Local anesthetic was injected subcutaneously along the incision site and the skull was prepared for incision. After exposing the cranial bone, the anterior/posterior axis was determined by positioning the instrument (injection needle) at ear bar zero. Six small craniotomies (roughly 0.5cm) were made using an air drill to expose the dura, and a 100µl Hamilton syringe fitted with a 25 gauge needle was mounted on the micromanipulator of the stereotaxic instrument. A suspension of genetically modified, replication-deficient, AAV (1e12 DNAse resistant particles/ml) that encodes eGFP or a fragment of human mutant huntingtin was injected through the 100µl Hamilton syringe connected to a Quintessential Stereotaxic Injector pump. The infusion rate began at 0.5µl/min then increased by 0.5 µl every 5 minutes. This progressive increase in infusion rate is known as convection enhanced delivery, and it improves the spread of infusate compared to conventional infusion methods^28,29^. The needle was left in place for an additional 5 min to allow the injectate to diffuse from the needle tip before removing. For the eGFP studies, monkeys received a total of 3 microinjections per hemisphere: one into the anterior head of the caudate nucleus (80µl), one into the anterior putamen (80µl) and a third in the post-commissural putamen (70µl). A total of 2.3E11 vector genomes were injected per hemisphere (see **Table 1** for summary of each surgical case). For the mHTT studies, we increased the number of injection sites in the caudate to improve spread within the structure such that these monkeys received 4 microinjections per hemisphere (anterior head of caudate, 80 µl; posterior head of caudate, 40 µl; anterior putamen, 95 µl; post-commissural putamen, 85 µl). A total of 3E11 vector genomes were injected per hemisphere. After microinjections were completed, the skull opening was filled with gelfoam, the incision closed, and the animal monitored closely during recovery. Post-operatively, animals were monitored for 5-7 days by veterinary staff and received Cefazolin, Hydromorphone, and Buprenorphine (antibiotic and pain management).

### NECROPSY/TISSUE COLLECTION

Necropsies and tissue collection were performed as previously described^52^. Briefly, animals were sedated with Ketamine and then deeply anesthetized with sodium pentobarbital followed by exsanguination. Brain and spinal cord were perfused with 2L of cold, sterile 0.9% saline. Brain was removed from the skull, placed into an ice-cold, steel rhesus macaque brain matrix and blocked into 4-mm-thick slabs in the coronal plane. Brain slabs were subsequently post fixed in 4% paraformaldehyde for 48 hours then cryoprotected in 30% sucrose for subsequent histological analyses.

### TISSUE PROCESSING/STAINING

Cryoprotected tissue was sectioned at 40 microns in the coronal plane on a freezing microtome. For immunohistochemical staining, sections were incubated with antibodies against eGFP (Invitrogen, A6455, 1:1000) or mHTT protein (1-82aa, Millipore, MAB5492, 1:500), and the appropriate biotinylated goat anti rabbit or goat anti mouse or secondary antibody (Vector Laboratories, BA-1000, BA-9200, 1:500). The signal was developed using a standard Vectastain ABC kit (Vector Laboratories, PK6100) with subsequent incubation in 3,3’-Diaminobenzidine (DAB) (Sigma, 112080050) and Nickel (II) Sulfate Hexahydrate (Sigma, N4882) intensification. For double labelling, sections were incubated overnight at room temperature with an antibody against mHTT protein (1-82 aa, Millipore, MAB5492 1:500), and either NeuN (Millipore, ABN78, 1:500) or GFAP (DAKO, 20334, 1:1000). The sections were then incubated for one hour with a goat anti mouse AlexaFluor 488 conjugated secondary antibody (Invitrogen, A11029, 1:500), and a goat anti rabbit AlexaFluor 546 conjugated secondary antibody (Invitrogen, A11035, 1:500) as previously described^56^.

### MICROSCOPY

20X Images for Figure 1, Figure 2B, and Figure 4 were taken on an Olympus BX51 microscope with an Olympus DP72 camera controlled by the CellSens program. Brightfield Images for Figure 2A were acquired on an Olympus VS120 Slide scanner utilizing an Olympus BX61VS microscope. An Allied Vision VC50 camera captured single Z-plane images through a 10x Objective. For double label fluorescence, a Leica SP5 confocal microscope with a 63x oil immersion objective was used along with the corresponding LAS AF program. The gain and exposure parameters were optimized for each confocal image.

### MEASURES OF VIRAL TRANSPORT/EXPRESSION

#### Cell counts

To characterize the biodistribution of AAV2 or AAV2.retro following injections into the striatum, we first identified regions of interest (ROIs) in the right hemisphere of the caudate (Cd), putamen (Put), amygdala (AMY), thalamus (THAL), dorsal prefrontal cortex (DPFC), dorsal premotor cortex (DPMC), anterior cingulate cortex (ACC), and somatosensory cortex (SSC). The contralateral hemisphere was used for molecular studies not described in the current manuscript. For each monkey, we collected three 10x IHC images stained for eGFP from 3 consecutive coronal sections from each of these ROIs (for a total of 9 images per region per animal). 10X images of caudate and putamen were placed adjacent to the sites of injection. The number of eGFP^+^ cells in each image was manually counted using ImageJ and averaged across the 9 images for each structure.

#### Ratings of relative staining density

We observed eGFP^+^ staining in many additional brain areas beyond the 8 ROIs in which cells were counted. We ranked the relative levels of transduction between brain areas to more fully characterize the biodistribution of AAV2.retro-eGFP. Brain regions, defined using the anatomical boundaries of Saleem & Logothetis^41^, were assigned the rank of high, mid, or low, depending on the relative level of transport based on the density of GFP+ cells in each ROI. The same ranking scheme was applied to coronal sections from animals injected with the AAV2.retro-85Q construct and stained with 1-82aa.

#### Area fraction fractionator

The Area Fraction Fractionator (Microbrightfield) was used to quantify the area fraction of eGFP positive cells in the putamen and caudate. Three 40-Micron thick coronal sections were chosen for each brain region to capture an anterior to posterior representation of the transduced regions. Coordinates of sections according to the atlas by Saleem and Logothetis were A-P: +28.0, +24.0, +21.0, and +17.0. Images captured at 10x on a VS120 slide scanner were uploaded to the Stereo Investigator program and the region of interest was outlined. One marker was used to select points that fell within the region of interest, and a second marker was used to select points containing eGFP-positive cells. The counting frame area was 1200 × 1200 μm, XY placement was 2000 × 2000 μm, and grid spacing was 50 μm. The area fraction estimation of eGFP+ cells in the caudate and putamen was determined by dividing the area of eGFP+ cells by the total area analyzed for each brain region.

### STATISTICAL ANALYSIS

All statistical analyses were performed using GraphPad Prism v8. To assess group differences in vector biodistribution, we first compared the numbers of GFP+ cells, or the area fraction of GFP+ cells, in each brain ROI using 2-way ANOVAs (Serotype x ROI). Subsequently, we ran planned comparisons between brain regions and serotypes using one-tailed, independent-sample t-tests for each comparison.

### 3D BRAIN VISUALIZATIONS

As part of ongoing studies in the lab, we created an MRI template from T2w SPACE scans collected from 18 healthy adult male and female rhesus macaques. Cortical and subcortical ROIs were manually defined on the template using ITK-SNAP^57^ according to the anatomical boundaries described by Saleem & Logothetis^41^. Next, ITK-SNAP was used to generate a 3-dimensional surface mesh for each ROI. The surface meshes were subsequently visualized with paraView 5.7.0 software^44^ and shaded according to the qualitative ranking of staining density: high-level transduced brain regions were shaded dark blue, mid-level regions were shaded royal blue, lower-level regions were shaded turquoise, regions with minimal to no staining were shaded gray, and injected ROIs (caudate and putamen) were shaded yellow.

## ACKNOWLEDGEMENTS

We thank Dr. Alla Karpova and colleagues at the Howard Hughes Medical Institute Janelia Research Campus for allowing the use of their pAAV2.retro-eGFP plasmid used in these studies. We thank Christoph Kahl and Michelle Gomes in the OHSU/ONPRC MVSC Core for production of all recombinant AAV viral vectors described here and for performing neutralizing antibody analyses. We thank the ONPRC Division of Comparative Medicine for the outstanding care of our rhesus macaques, with special thanks to Brandy Dozier, Lauren Drew Martin and Theodore Hobbs. We thank the ONPRC Pathology Unit for their expertise and assistance with necropsy and pathology. This research was supported by NIH/NINDS Award NS099136 (J.L.M), NIH/NINDS Award NS110149 (A.R.W), and a generous donation from Quentin and Bee Neufeld (J.L.M). This work was also supported by NIH Core Grant P51OD011092 and the NIH Instrumentation Grant 1S10OD025002-01.

## AUTHOR CONTRIBUTIONS

**Study design:** A.R.W., W.A.L., J.L.M.

**Surgical delivery of AAVs**: A.R.W., W.A.L., D.B., J.S.D., J.L.M.

**Necropsy and tissue collection:** A.R.W., W.A.L., D.B., J.S.D., J.L.M.

**Histology and microscopy:** W.A.L., A.R.W., J.L.M.

**Statistical analysis:** A.R.W, W.A.L., J.L.M.

**Figure preparation:** A.R.W, W.A.L., J.L.M.

**Manuscript preparation:** A.R.W, W.A.L, J.L.M.

All authors reviewed the manuscript.

## ADDITIONAL INFORMATION

### Competing Interests

J.L.M. has previously been a paid consultant for Spark Therapeutics and is currently a paid consultant/key opinion leader for Takeda Pharmaceutical Company. J.L.M. is also currently receiving financial compensation for consulting with investigators at Rush University Medical Center and The University at Albany-SUNY regarding an award funded via the Michael J. Fox Foundation for Parkinson’s Research. All other authors have no conflicts to report.

### Data Availability

The authors will make all materials, data and associated protocols generated as part of this publication available to readers upon request.

