## Supplemental material for "Intra-striatal AAV2.retro administration leads to extensive retrograde transport in the rhesus macaque brain: implications for disease modeling and therapeutic development"

**Supplemental Table S1.**

| **AAV2.retro** | | | | | | | | | |
|---|---|---|---|---|---|---|---|---|---|
| | Caudate | Putamen | ACC | AMY | DPFC | DPMC | PMC | SSC | THALAMUS |
| Caudate | | 0.0003 | 0.0051 | 0.0011 | 0.0006 | 0.0009 | 0.0003 | 0.0002 | 0.0009 |
| Putamen | 0.0003 | | <0.0001 | <0.0001 | <0.0001 | <0.0001 | <0.0001 | <0.0001 | <0.0001 |
| ACC | 0.0051 | <0.0001 | | 1.0000 | 1.0000 | 1.0000 | 0.6605 | 0.2904 | 1.0000 |
| AMY | 0.0011 | <0.0001 | 1.0000 | | 1.0000 | 1.0000 | 1.0000 | 1.0000 | 1.0000 |
| DPFC | 0.0006 | <0.0001 | 1.0000 | 1.0000 | | 1.0000 | 1.0000 | 1.0000 | 1.0000 |
| DPMC | 0.0009 | <0.0001 | 1.0000 | 1.0000 | 1.0000 | | 1.0000 | 1.0000 | 1.0000 |
| PMC | 0.0003 | <0.0001 | 0.6605 | 1.0000 | 1.0000 | 1.0000 | | 1.0000 | 1.0000 |
| SSC | 0.0002 | <0.0001 | 0.2904 | 1.0000 | 1.0000 | 1.0000 | 1.0000 | | 1.0000 |
| THALAMUS | 0.0009 | <0.0001 | 1.0000 | 1.0000 | 1.0000 | 1.0000 | 1.0000 | 1.0000 | |

| **AAV2** | | | | | | | | | |
|---|---|---|---|---|---|---|---|---|---|
| | Caudate | Putamen | ACC | AMY | DPFC | DPMC | PMC | SSC | THALAMUS |
| Caudate | | 0.0012 | 0.0141 | 0.0140 | 0.0140 | 0.0139 | 0.0141 | 0.0139 | 0.0141 |
| Putamen | 0.0012 | | <0.0001 | <0.0001 | <0.0001 | <0.0001 | <0.0001 | <0.0001 | <0.0001 |
| ACC | 0.0141 | <0.0001 | | 1.0000 | 1.0000 | 1.0000 | 1.0000 | 1.0000 | 1.0000 |
| AMY | 0.0140 | <0.0001 | 1.0000 | | 1.0000 | 1.0000 | 1.0000 | 1.0000 | 1.0000 |
| DPFC | 0.0140 | <0.0001 | 1.0000 | 1.0000 | | 1.0000 | 1.0000 | 1.0000 | 1.0000 |
| DPMC | 0.0139 | <0.0001 | 1.0000 | 1.0000 | 1.0000 | | 1.0000 | 1.0000 | 1.0000 |
| PMC | 0.0141 | <0.0001 | 1.0000 | 1.0000 | 1.0000 | 1.0000 | | 1.0000 | 1.0000 |
| SSC | 0.0139 | <0.0001 | 1.0000 | 1.0000 | 1.0000 | 1.0000 | 1.0000 | | 1.0000 |
| THALAMUS | 0.0141 | <0.0001 | 1.0000 | 1.0000 | 1.0000 | 1.0000 | 1.0000 | 1.0000 | |

**Supplemental Table S1. Matrix of p-values from post-hoc comparisons of cell counts between brain regions.** Tables report the p-values from one-tailed independent sample t-tests to compare the numbers of GFP+ cells in each ROI separately for each of the two serotypes, AAV2.retro (upper panel) and AAV2 (lower panel).

**Supplemental Table S2.**

| High | Medium | Low | None |
|---|---|---|---|
| Dorsal prefrontal cortex | Somatosensory cortex | Parietal cortex | Occipital cortex |
| Ventral prefrontal cortex | Orbitofrontal cortex | Superior temporal cortex | Cerebellum |
| Dorsal premotor cortex | Rhinal cortex | Thalamus (MD/LD) | Thalamus (RE) |
| Ventral premotor cortex | Thalamus (LP/VPL) | Inferior temporal cortex | Thalamus (VA/VL) |
| Anterior cingulate cortex | Amygdala (BLN) | Substantia nigra, pars comp. | |
| Subthalamic nucleus | Claustrum | Globus Pallidus, int. segment | |
| Pre-supplemental motor cortex | Globus Pallidus, ext. segment | | |
| Supplemental motor cortex | | | |
| Primary motor cortex | | | |
| Insular Cortex | | | |

**Supplemental Table S2. Relative levels of brain-wide mutant HTT expression following intra-caudate and intra-putamen injection of AAV2.retro-HTT85Q**. Brain regions were qualitatively ranked relative to one another, considering cell number and density, and placed into categories of high, medium, low and minimal/none. All regions listed here had less expression compared to the injected regions of the caudate and putamen. MD- medial dorsal, LD- lateral dorsal, RE- nucleus reuniens, BLN- basolateral nucleus, int.- internal, ext.- external, VA- ventral anterior, VL- ventral lateral, LP- lateral posterior, VPL- ventral posterolateral.
